# Transient beta modulates decision thresholds during human action-stopping

**DOI:** 10.1101/2021.07.05.447605

**Authors:** Vignesh Muralidharan, Adam R Aron, Robert Schmidt

## Abstract

Action-stopping in humans involves bursts of beta oscillations in prefrontal-basal ganglia regions. To determine the functional role of these beta bursts we took advantage of the Race Model framework describing action-stopping. We incorporated beta bursts in three race model variants, each implementing a different functional contribution of beta to action-stopping. In these variants, we hypothesized that a transient increase in beta could 1) modulate decision thresholds, 2) change stop accumulation rates, or 3) promote the interaction between the Stop and the Go process. We then tested the model predictions using EEG recordings in humans performing a Stop-signal task. We found that the model variant in which beta increased decision thresholds for a brief period of time best explained the empirical data. The model parameters fitted to the empirical data indicated that beta bursts involve a stronger decision threshold modulation for the Go process than for the Stop process. This suggests that prefrontal beta influences stopping by temporarily holding the response from execution. Our study further suggests that human action-stopping could be multi-staged with the beta acting as a pause, increasing the response threshold for the Stop process to modulate behavior successfully. Our novel approach of introducing transient oscillations into the race model and testing against human neurophysiological data allowed us to discover potential mechanisms of prefrontal beta, possibly generalizing its role in situations requiring executive control over actions.

## INTRODUCTION

Prefrontal beta oscillations occur during action-stopping in both human and non-human primates (Errington et al., 2020; Hannah et al., 2020; Jana et al., 2020; Swann et al., 2009; Wagner et al., 2018). The Stop-signal task is widely used to assess action-stopping behavior in humans. The task mostly involves Go trials, in which subjects quickly respond to a Go cue. However, occasionally, on Stop trials, a Stop-signal is presented after the Go cue, instructing the subject to withhold responding. There are increases in prefrontal beta in these Stop trials, and these increases happen before the action is stopped, i.e. before the stop-signal reaction time (SSRT). It has been hypothesized that these beta oscillations (which occur as bursts at the single-trial level) could be a marker of a fast hyper-direct prefrontal STN pathway that gets recruited to stop motor-processes (Aron, 2011; Chen et al., 2020). Furthermore, the timing of beta bursts correlates with the time the action is cancelled (Hannah *et al.*, 2020; Jana *et al.*, 2020). Currently, the neural mechanisms by which prefrontal beta influences action-stopping are not clear. One possibility is that a beta burst reflects an active communication channel between prefrontal regions and the basal ganglia (c.f. (Fries, 2005)), e.g. as a form of top-down control biasing the current behavioral strategy towards stopping via STN to inhibit motor processes (Schmidt et al. (2019)). However, as there are other possibilities as well, it still remains unclear how beta relates to action-stopping.

In the well-developed race model framework, a Go and a Stop process race against each other and whichever process reaches the decision threshold first determines the behavioral outcome (Go or Stop) (Logan and Cowan, 1984; Logan et al., 2014; Verbruggen and Logan, 2009). Here we developed and studied three different race model variants of how beta could modulate the Go and Stop processes by affecting decision thresholds, accumulation rates, and the interaction between Go and Stop processes, respectively. For each model variant, we derived predictions for the relationship between beta and behavioral data, and then tested these predictions in two data sets of EEG recordings in humans performing a stop-signal task. We found that the human data is best described by a race model in which beta bursts transiently affect the decision threshold, providing a new functional role of prefrontal beta in human action-stopping.

## METHODS

### Dataset/Participants

We analyzed data from existing data-sets. One was a reanalysis, (Dataset-1, N = 13, Jana et al. 2020; Mean Age = 20 ± 0.5 years, eight females, all right-handed), and the other was unpublished (Dataset-2: N = 26, Mean Age = 21 ± 0.5 years, 16 females, all right-handed except one participant who was left-handed). All participants provided written informed consent according to a UCSD Institutional Review Board protocol and were compensated at $20/hr. One participant was removed from analysis in Dataset-1 due to misalignment of EEG markers and behavior. Three participants were removed in Dataset-2, two participants had noisy EEG data and one had estimated Stop-signal reaction time of <100ms. Thus, the final sample size was N = 12 in Dataset-1 and N = 23 in Dataset-2.

### Stop-Signal Task

Both datasets were acquired with the behavioral task run using MATLAB 2014b (Mathworks, USA) and Psychtoolbox (Brainard, 1997). The task was a visual stop-signal task where a trial began with a white square at the center of the screen for 500 ± 50ms. Following this a right or left white arrow appeared at the center. Participants pressed a button with either their right index finger, when a left arrow appeared, or their right pinky (Dataset-1) or middle (Dataset-2) finger when a right arrow appeared. They were instructed to do this as fast and as accurately as possible (Go trials). The stimuli remained on the screen for 1 s. A warning ‘Too Slow’ was presented if the participants did not make a response within this time and the trial was aborted. On a minority of trials (25%), the arrow turned red after a Stop Signal Delay (SSD), and participants tried to stop the response (Stop trials). The SSD was adjusted using two independent staircases (for right and left directions), where the SSD increased and decreased by 50ms following a Successful Stop and Failed Stop, respectively. Each trial was followed by an inter trial interval and the entire duration of each trial including the inter trial interval was 2.5 s. There were in total 1920 trials (1440 Go trials and 480 Stop trials) and 400 trials (300 Go trials and 100 Stop trials) per participant in Dataset-1 and Dataset-2 respectively.

### Data analysis & computational modelling

All analyses and simulations were performed using MATLAB R2016b.

#### Electroencephalography (EEG)

EEG data were recorded using 64 channel scalp EEG in the standard 10/20 configuration using an Easycap system (Easycap and BrainVision actiCHamp amplifier, Brain Products Gmbh, Gilching, Germany) for Dataset-1 and the ActiveTwo system (Biosemi Instrumentation, The Netherlands) for Dataset-2. The EEG signals were digitized at 1024 Hz and pre-processed using EEGLAB13 (Delorme and Makeig, 2004) and custom-made MATLAB scripts. The data were downsampled to 512 Hz and band-pass filtered between 2–100 Hz. A 60, 120 and 180 Hz FIR notch filter were applied to remove line noise and its harmonics. EEG data were then re-referenced to the average. The continuous data were visually inspected to remove bad channels and noisy stretches.

To look at prefrontal (or right frontal) beta, we used the ICA method and Time-Frequency analysis to obtain a putative right frontal spatial filter (done exactly as in Jana et al. (2020)). After rejecting non-brain related independent components (ICs), identified from the frequency spectrum (increased power at high frequencies), scalp maps (activity outside the brain) and the residual variance of the dipole (greater than 15%), we selected a putative right frontal IC from the scalp maps (if not present then we used frontal topography). The channel data were then projected onto the corresponding right frontal IC. The right frontal IC was validated by evaluating the time-frequency plots for successful stop trials and confirming a beta power increase (13-30 Hz) between Stop-signal and Stop-signal reaction time (SSRT) in Successful Stop trials. To do so, we first epoched the data from −1500 to 1500ms for all trials type: Successful, Failed Stops and Correct Go trials (in relation to Stop-signal in Stop trials; and in relation to Go-cue in Correct Go trials). We then used Morlet wavelets for computing the time-frequency plots (4-30Hz) in Successful Stop trials, with 3 cycles at low frequencies and linearly increasing by 0.5 for higher frequencies. The beta frequency having the maximal power within Stop-signal and SSRT in these trials was also estimated for each participant as their peak beta frequency.

#### Beta burst extraction

The beta burst extraction was also done as in Jana et al. (2020), which was adapted from Little et al. (2019). The epoched data were filtered at the peak beta frequency for each participant using a Gaussian window with full-width half maximum of 5Hz. From the resulting complex analytic time-series, we obtained the power estimate by computing the absolute of the Hilbert transform of this time-series. In each participant, to define the burst threshold, the beta amplitude within a period of −1000 to −500 ms (i.e. prior to Stop-signal in the Stop trials, and prior to mean SSD in the Correct Go trials) was pooled across all trials. The threshold was set as the median + 1.5 SD of the beta amplitude distribution. Once the burst was detected, the burst width threshold was set as the median + 1 SD. Burst % was computed by binary-coding the time-points where the beta amplitude crossed the burst width-threshold. For each detected burst, the time of the peak beta amplitude was marked as the burst time. We also computed the times at which a beta burst ended, i.e. the beta amplitude fell beyond the burst width threshold and marked it as the burst offset time.

#### Analyzing behavior in relation to beta bursts in Stop trials

To analyze behavior in the Stop-signal task we obtained Go and Failed Stop reaction times. SSRTs were estimated using the integration method (Verbruggen et al., 2019). Using the extracted timing of the beta bursts (see above), we estimated the probability of responding to the Stop cue, P(Respond), as a function of the time relative to the burst. To obtain reliable estimates this was done across participants, as a fixed-effects analysis, considering the behavior of the population as one. We pooled all Stop trials with at least one beta burst between −100ms and the corresponding SSRT (in relation to Stop-signal). From the pooled data, we then included Stop trials from those SSDs for which we had at least 50 trials. We did so as this was a good tradeoff between having enough trials to estimate P(Respond) reliably and to eliminate the really short and long SSDs where firstly there were not many trials and secondly where the effect of beta burst time as predicted by the modelling analyses was the least (see Fig. 1d-f). Trials with more than one burst in the selected time-window were split into several trials, each with a single corresponding burst time. P(Respond) was then estimated at different time points relative to the time of the Stop-signal with a moving 50ms-wide window centered ranging from −50ms to 250ms in steps of 1ms. P(Respond) was then simply taken as the fraction of failed stop trials in that 50ms window. To determine whether a given P(Respond) significantly differed from chance level we used a permutation test (with 1000 permutations), in which the labels of Successful and Failed Stop trials were shuffled to yield surrogate P(Respond) distributions. Time points in which the empirical P(Respond) was smaller than (1-0.05/n)*100% of the surrogate values were considered significant at a p-value threshold of 0.05 with a Bonferroni correction for n= 5 multiple comparisons (given that there are 5 non-overlapping windows in our time period of interest, i.e. 0 and 250ms).

**Figure 1.**
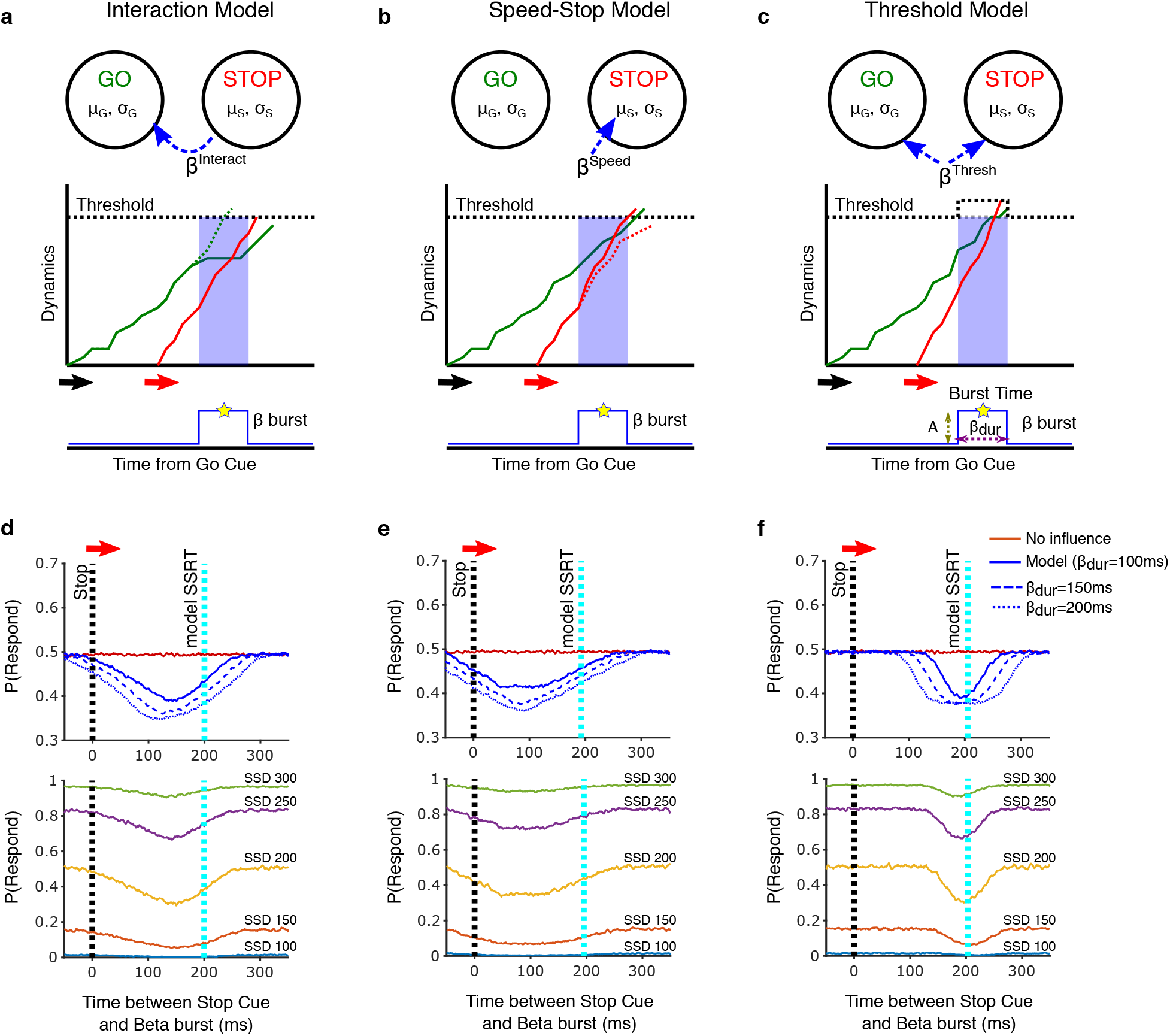
Race Model dynamics with transient bursts. Schematic of the model setup and the accumulation dynamics of the Go and Stop processes for the three race model variants: a) Interaction Model, b) Speed-Stop Model and c) Threshold model. The solid green and red lines represents the Go-process and Stop-process accumulation dynamics, respectively. The dotted green and red lines represent the dynamics of the Go and Stop-process without the presence of a beta burst for comparison. The dotted black line shows the threshold, which in case of the Threshold model increases whenever the burst is present. A beta burst (solid blue line) is parametrized by the time it occurs (Burst Time), its duration (β_dur_) and its amplitude (A). d-f) Corresponding predictions of each of the three model variants on how a beta burst after the stop cue affects the probability of the Go process to win the race (i.e. P(Respond)). For each model variant (arranged as in panel a, b and c), we show the probability to respond (P(Respond)) for varying time intervals between Stop cue and the center of the beta burst (x-axis). Longer beta burst durations (β_dur_ = 100, 150 and 200ms) prolonged and strengthened the effect on P(Respond). We chose model parameters here to yield RTs and SSRTs in a range that is typical for this task (see Methods), and illustrate in the plots below the effect of different SSDs (for β_dur_ = 100). The dotted black and cyan lines represent the time of the Stop cue and model SSRT respectively.

#### Analyzing behavior in relation to beta bursts in Go trials

The relationship between beta bursts and Go RTs was examined by estimating the probability distribution of burst offset times for different Go RTs. As above, for the estimation of P(Respond), this required a large number of trials, so data was again pooled across participants. Here all correct Go trials with at least one beta burst occurring in the time window between - 100ms relative to the Go cue and that trial’s Go RT were included. As before, trials with more than one burst were split into multiple trials with the same Go RT but different beta burst times. We decided to look at RTs which fell in the range of 300-500ms (for Dataset-1) and 300-700ms (for Dataset-2) as they constituted the majority of the RT distribution across individuals. For each 50ms-wide Go RT bin we looked at the probability (or fraction) of the burst offset times occurring at each time-point between −50 to 500ms in relation to the Go cue (−50 to 700ms in case of Dataset-2). This produced a probability distribution map of Go RTs and burst offset times. We also quantified this data in a different way by looking at the distribution of the difference between all Go RTs (not just the range selected for the probability map) and the burst offsets. We compared this to a null distribution, in which we assumed that the burst offset times were equiprobable at all times between −100ms to Go RT+100ms.

#### The three race model variants

We considered three race model variants describing the influence of beta on behavior. Each variant followed the standard race model differential equations governing the accumulation dynamics for both the Go and Stop decision-variables (Boucher et al., (2007)):

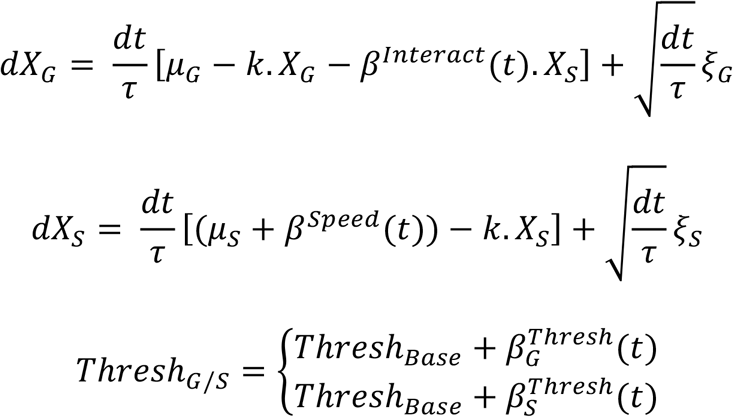

In our simulations dt/τ was set as 0.001. The leakage term k for both processes was set to 0. μ_G_ and μ_S_ parameters are the average accumulation rate for the Go and Stop-process respectively, ξ_G_ and ξ_S_ are random variables representing the stochastic accumulation dynamics for the Go and Stop-process respectively, drawn from a normal distribution N(0, σ_G/S_). Here σ_G_ and σ_S_ represent the standard deviation of the normally distributed noise term for the Go and Stop-process respectively. The baseline decision thresholds for both the Go and Stop processes were set to 1 (Thresh_Base_). For the three different versions of the model (Interaction, Speed-Stop and Threshold models), we added a time-varying β term (β^Interact^, β^Speed^, β^Thresh^) to introduce the influence of a beta burst in the corresponding model scenario. The beta burst was modelled as a step function with pulse duration as the length of the burst (β_dur_), where the center of the pulse corresponded to the simulated burst time (also see Fig. 1).

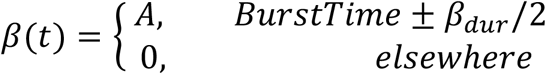

Here A is the amplitude/strength of beta burst. In each model variant, the respective beta variable was varied whilst the other two beta parameters were set to zero. Usually, the outcome of the race was given by process reaching the threshold first. However, in the Threshold model, there were trials, in which the reset of threshold to the baseline value (i.e. when the beta burst ended) lead to both the Go and Stop processes being above the threshold. In such cases the process which had the larger activation was considered as the winner of the race. The model Go reaction times (Go RTs) and Stop-signal reaction times (or model SSRT) were estimated as the time the Go and Stop-process took to reach the threshold, respectively.

#### Model parameters

For the initial examination of the three race model variants, we used parameter settings that produced typical behavior data seen in Stop-signal tasks. By choosing the Go and Stop-process parameters to be μ_G_ = 2.5, σ_G_ = 0.006 and μ_S_ = 5, σ_S_ = 0.006 respectively, we obtained simulated Go RTs of ~400ms and SSRTs of ~200ms. In each trial of the model, the SSD was randomly selected from a range of 100-300ms with 50ms resolution. We then determined P(Respond) as a function of when a beta burst occurred similar to the method described above for the empirical data. However, in the models we could systematically vary the time of the beta bursts, and examined a range from −100ms to 250ms relative to the Stop-signal in 1ms steps. We then computed P(Respond) for a particular burst time as the number of failed stop trials divided by the total number of stop trials for that particular SSD. We also varied the duration of the beta burst by setting the β_dur_ parameter to 100, 150 and 200ms in different simulations.

After the general, initial examination of the three model variants (Fig. 1), the Threshold model was also fitted to individual participant data (Fig. 2). For each iteration of the fitting procedure 30,000 trials (15,000 Go trials and 15,000 Stop trials) were simulated. Since the incidence of beta bursts is generally low during stopping, ~20% (Errington *et al.*, 2020; Hannah *et al.*, 2020; Jana *et al.*, 2020; Wessel, 2020), we simulated a beta burst in only 20% of all trials (total 6000 trials, 3000 Go and 3000 Stop trials). In each trial with a burst the time of the beta burst was drawn as a random time point in between the Go cue and the slowest RT of the corresponding participant with a resolution of 100 possible time points. The fitting then proceeded in two stages, starting with the parameters of the Go process (μ_G_, σ_G_ and A_G_). The squared difference between experimental and simulated correct Go RT CDFs was used as the optimization function. In the second stage, we then fixed the parameters of the Go process to the best fit, and then fitted the stop parameters (μ_S_, σ_S_ and A_S_), using the squared difference between experimental and simulated inhibition functions as the optimization function. For the experimental inhibition functions, we first fitted a cumulative Weibull function, which best captures the shape of inhibition function (Hanes et al., 1998), and then used that as the cost function for our parameter optimization. In each of the two stages of the fitting procedure initially a coarse grid search was performed, in which we provided a range of parameters for μ, σ and α and determined the top 20 best fits. These were then used as initial conditions for the Nelder-Mead Simplex algorithm (*fminsearch* function in MATLAB) with maximum function evaluation of 600 iterations. We compared the findings to a null model scenario where there was no threshold modulation, i.e. simulating trials by setting the amplitude of the threshold parameter for both the go and stop-process as zero (A_G/S_ = 0).

**Figure 2.**
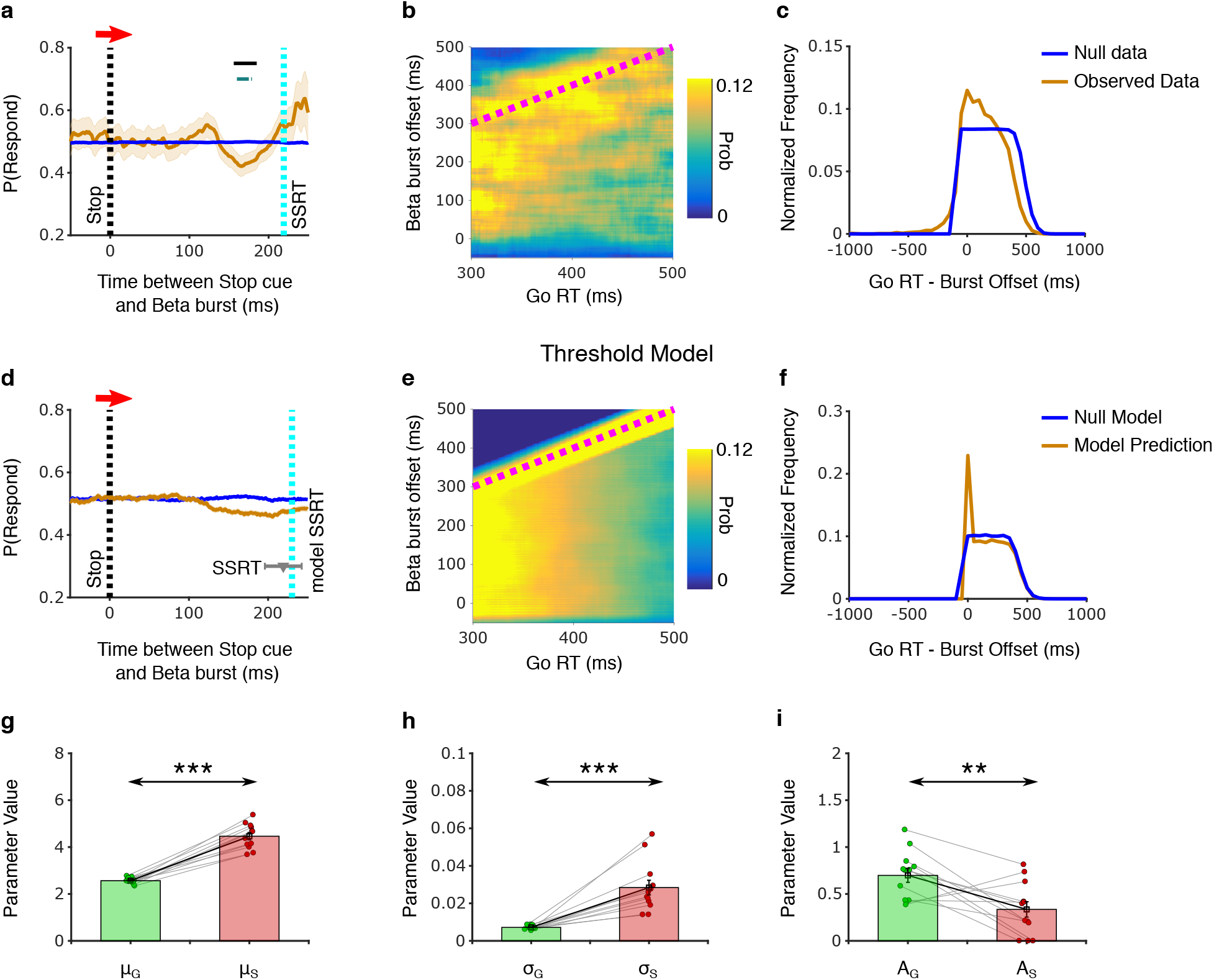
Beta bursts modulate decision thresholds. a) Probability of response as a function of time between Stop cue and beta burst in experimental data for Dataset-1 (see Methods). Horizontal lines indicate time points with significant differences between the observed and shuffled data (black line shows p<0.05; blue line with Bonferroni correction). b) The joint probability distribution between the timing of beta burst offsets and Go RTs. The diagonal dotted magenta line indicates when the Go RT is equal to the burst offset time. c) The normalized distribution of the difference between Go RTs and burst offset times, compared to a null-distribution (solid blue line). d-f) Same as (a-c) but for the Threshold model. In d) the solid blue line represents the outcome of the model without any threshold modulation for comparison. The grey triangle and error bar represents the mean and standard deviation of the SSRT from experimental data; the model SSRT lies within the standard deviation of the experimental data. g-i) Fitted mean slope μ (g), slope standard deviation σ (h), and threshold modulation A (i) parameters for go and stop-process.

## RESULTS

### Three models of beta in stopping

We considered three different models for how a beta burst at a specific time (burst time) could modulate the race between the Go and the Stop process. Firstly, in the Interaction model (Boucher *et al.*, 2007) the Stop-process inhibited the Go-process during a beta burst for a duration β_Dur_ (Fig. 1a). This would slow the rate of accumulation in the Go-process, making it more likely for the Stop-process to win the race. Secondly, in the Speed-Stop model the occurrence of a beta burst increased the rate of accumulation in the Stop-process (Fig. 1b), making it more likely for it to reach decision threshold first. Finally, in the Threshold model a beta burst would increase the decision threshold of both the Go and Stop processes (Fig. 1c). Since the rate of accumulation is generally quicker in the Stop-process than in the Go-process, the threshold increase would buy more time for the Stop-process to reach the threshold first and win the race.

While all three models implemented the hypothesis that beta decreases the probability of responding, we examined whether the predicted time course and extent of the modulation differed across models. Here, and in the analysis of the experimental data below, we looked at the probability of responding P(Respond) as a function of the time interval between the beta burst and the Stop-signal. For example, a P(Respond) of 0.6 at 0.3s would mean that 60% of the trials with a beta burst centered at 0.3s after stop cue were failed stop trials. We simulated each model by choosing parameters for the Go-process (μ_G_ = 2.5, σ_G_ = 0.006) and the Stop-process (μ_S_ = 5, σ_S_ = 0.006) that yielded mean go RTs of ~400ms and SSRTs of ~200ms, which are in the range of typical values for this task. We found that the pattern of modulation had a U-shaped profile in each of the three models, with larger amplitudes for longer burst durations (Fig. 1d-f). Furthermore, the modulation was strongest for intermediate SSDs (200ms) compared to longer or shorter ones. However, the fine time course and the relation to the SSRT, differed across the three models. For the Interaction model and the Speed-Stop model the trough of the U-profile was seen well before the model SSRT. In contrast, in the Threshold model the modulation pattern was very different, with a sharper modulation in a narrower time window just before the model SSRT. Next, we then compared these model predictions with experimental data to see whether beta affects behavior in a similar way, and whether beta modulation can be best described by one of these model variants.

### Beta bursts affect going and stopping by modulating decision thresholds

To test the model predictions, we employed two data sets of humans performing stop-signal tasks with simultaneously recorded EEG (see Methods). The behavioral data showed a pattern that is typical for stop-signal tasks with Go RTs being longer than Failed Stop RTs (Dataset-1: 406 ± 6ms vs 373 ± 6ms, t_1,11_ = 14.7, p < 0.001; Dataset-2: 468 ± 16ms vs 411 ± 14ms, t_1,22_ = 15.8, p < 0.001). The P(Respond) was also nearly 50% (Dataset-1: 49.6 ± 0.2%; Dataset-2: 51.7 ± 0.7%) suggesting that the staircase procedure worked well. The SSRT computed via the integration method was 219 ± 7ms and 222 ± 6ms for Dataset-1 and Dataset-2 respectively.

We first examined in the experimental data whether stopping changes as a function of the time interval between the beta burst and the Stop-signal. To do so, we selected the subset of trials in which there was at least one beta burst in the time between 100ms before the stop cue and the SSRT (see Methods). Thereby, each trial provided a time point (given by the interval between the stop cue and the time of peak of the beta burst) and the outcome (respond or not). Pooling these trials over participants allowed us then to estimate the probability of responding, P(Respond), at each time point, as we did in our model investigations (Fig. 1d-f). In the experimental data, we found that P(Respond) was unchanged for beta bursts briefly after the stop cue, but then decreased closer towards the SSRT (Fig. 2a). To determine whether this modulation in P(Respond) is a significant deviation from chance, we compared it to a null-distribution estimated from shuffling the labels of successful and failed stop trials. We found a rather narrow time window with a significant modulation ~40-60ms before the SSRT (Fig. 2a). While the narrow time window of the modulation seemed to match the Threshold model, the timing of the peak modulation was also in the range suggested by the Interaction model (Fig. 1d). Furthermore, fitting the Threshold model parameters to the behavioral data in Dataset-1 (using β_dur_ of 100ms) also generated a wider modulation window (Fig. 2d), similar to the other models (Fig. 1a-c). Therefore, we concluded that examining the beta modulation of P(Respond) is by itself not sufficient to distinguish between the three models (also see Supplementary Fig. S2a for the P(Respond) for other burst durations).

As an alternative way to test the different models, we made use of the fact that a unique aspect of the Threshold model is that it also predicts an effect of beta in Go trials. The other two models (Interaction and Speed-Stop) predict that there would be no influence of beta in Go trials because the Stop-process is not activated (or only very weak). Therefore, to distinguish the models further, we then examined a possible beta modulation of Go trials.

We first established the relationship between beta and Go RTs in the Threshold model. As the beta duration varied, we focused here on the offset of beta, rather than using the center of a beta burst as above. We found that in the Threshold model the beta-RT relation was governed by an intricate pattern (Fig. 2e). For short reaction times (~300ms) there was a rather uniform distribution of beta in the relevant time window from 0 to 300ms relative to the Go cue. In contrast, for longer RTs, the distribution of the beta time points became more skewed, with more beta offset time points occurring around the RT. This was visible as a diagonal stripe with high beta offset probabilities (Fig. 2e).

This modulation pattern matched the intuition behind the Threshold model. For short RTs, the Go process is steep, and therefore a threshold modulation would not have a large effect at any time (thus the uniform distribution in Fig. 2e). However, for longer RTs, the Go process is less steep, and it would thus be more likely that the threshold modulation prolongs the RT. Furthermore, beta would not affect the RT if it occurred briefly after the Go cue because then the threshold modulation would have already ended by the time the Go process is close to the threshold. Instead, the beta offset would affect the RT if it occurred briefly before the RT, as a sudden drop in the threshold would then lead to a threshold crossing. We concluded that this intricate pattern of beta modulation in the Threshold model provided more specific predictions than the straightforward modulation of P(Respond).

Next, we tested the predictions of the Threshold model by looking at the correct Go trials pooled across all participants. As above we included only trials in which there was at least one beta burst in the time between the go cue and the response, and for each trial we estimated the time of the offset of the beta burst (see Methods). This allowed us to visualize the relation between Go RT and beta burst offset for the experimental data in the same way as we did for the Threshold model (Fig. 2b,e). Interestingly, for different Go RTs, there was a different distribution of the probability of beta burst offsets. For shorter RTs, beta burst offsets were approximately uniformly distributed over time. For longer RTs, the distribution became more skewed, with more beta offsets towards the corresponding RTs, yielding a diagonal stripe in the probability distribution (Fig. 2b). Even though this stripe was somewhat broader than in the Threshold model (see Discussion and also Supplementary Information Fig. S4), overall there was a striking resemblance to the quite specific predictions of how beta should affect Go RTs in the Threshold model.

One interpretation of this pattern of beta modulation is that responses are harder to execute as long as a beta burst is present, and once it ends, or is close to ending, the response emerges. To have a closer look at this relation, we represented the same data as the distribution of the time interval between the Go RT and beta offset in each trial. In line with our interpretation above, we found that the distribution peaked at small positive values, meaning that in most trials responses occurred briefly after the offset of the beta burst. We compared this distribution to a null distribution, which was based on the assumption that beta burst offsets were uniformly distributed (i.e. beta does not affect RTs). We found that our data was significantly different from the null distribution, with more burst-offset-times close to the RTs (Fig. 2c, Kolmogorov-Smirnov test, KSstat = 0.15; p < 0.001). Furthermore, we confirmed that the Threshold model exhibited a similar distribution (Fig. 2f), with a sharper peak reflecting the narrow diagonal stripe described above. To confirm the validity of our findings on the P(Respond) and on Go trials, we ran the same analyses on an independent dataset (Study 2, N = 23) and observed the same effects (Supplementary Information, Fig. S1). Overall, the analyses relating the offset of beta bursts with Go RTs provide evidence that beta bursts modulate decision thresholds.

Finally, we examined the parameters that were obtained for the Threshold model fitted to the experimental data (Fig. 2d-f; see Methods). Both mean and standard deviation of the Stop process were significantly larger than those of the Go process (Fig. 2g, h). Furthermore, the beta modulation of the threshold was on average higher for the Go process than for the Stop process (Fig. 2i), in line with the intuition that the increased threshold buys more time for the Stop process to overtake the Go process. However, inspection of the fitted values for the modulation of the Stop process (A_S_) indicated differences across participants. For the majority of participants there was no modulation of the Stop threshold at all (in line with the intuition) or lower than the Go process, while for some participants the Stop threshold modulation was of similar magnitude as the Go process threshold modulation. However, given that the Stop process slopes were consistently steeper than the Go process slopes, even a similar modulation of Go and Stop thresholds would effectively increase the probability of the Stop process to win the race. While in our default simulations beta bursts were only present in 20% of the trials (similar to the experimental data), we further confirmed these findings in a model variant in which beta bursts were present in all trials (see Supplementary Information, Fig. S3).

### Threshold modulation explains relationship between burst-time and stopping time

Building on the evidence that beta bursts modulated decision thresholds, we next tested whether the Threshold model also accounts for further, single-trial properties of stopping. In recent work, we demonstrated that beta bursts are linked to single-trial SSRTs measured via EMG (Hannah *et al.*, 2020). We examined this relation in successful stop trials, in which a beta burst occurred between the stop cue and the corresponding single-trial SSRT. Pooling data across participants, we determined the distribution of the time intervals between beta burst and single-trial SSRT, and found a peak in the histogram when the beta burst preceded correct stopping by 50ms (Fig. 3a). This was in comparison to a null distribution, in which we assumed that a beta burst would occur randomly (uniformly distributed) anytime between the Stop cue and the trial’s SSRT. Applying the same analysis in the Threshold model, we saw that the distribution of time intervals between beta burst and single-trial SSRT was very similar. As in the experimental data there was a peak at 50ms in the model. The peak was even sharper in the model, probably due to the simplified, noise-free composition of the model (also see Supplementary Information, Fig. S4). The relation between beta and stopping occurred in the model even though the threshold modulation primarily affected the Go process. However, in successful stop trials a beta burst occurred with an increased probability ~50ms before stopping because in this time window the increase of the Go threshold affects the outcome of the race most. We conclude that the Threshold model accounts for several aspects of behavioral and electrophysiological data in both Go and Stop trials.

**Fig. 3.**
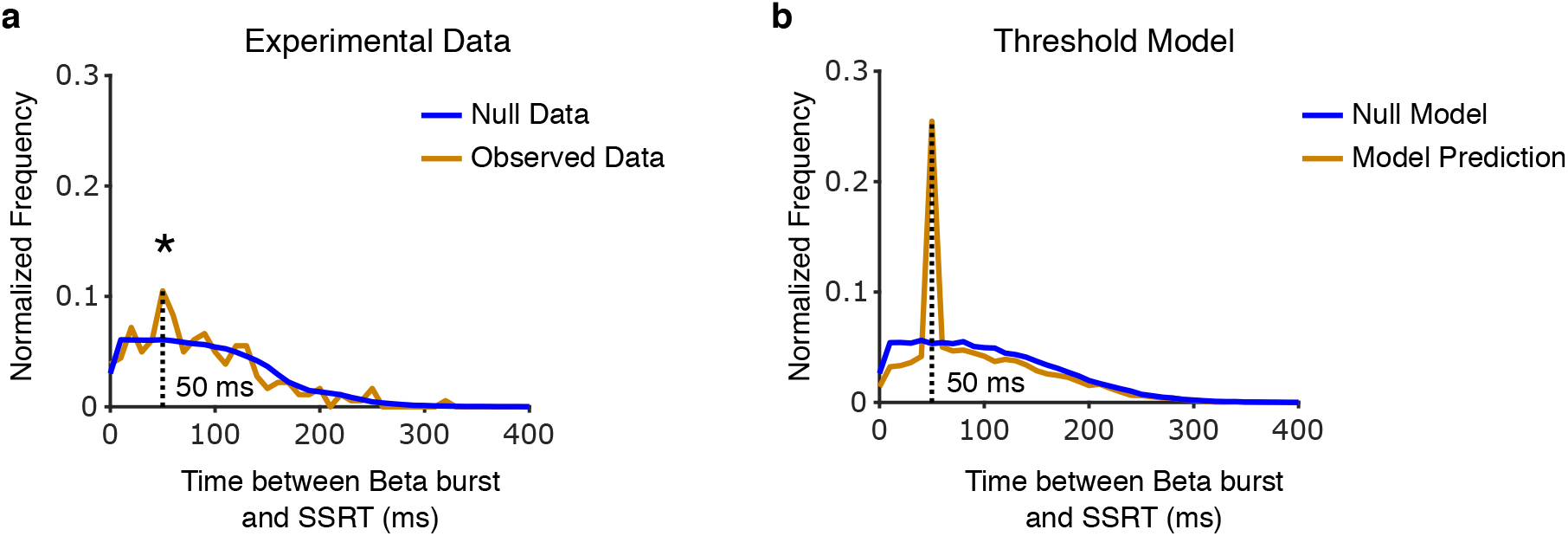
Threshold modulation is most effective when it is close to the SSRT. The distribution of the difference between the single-trial SSRT and the beta burst in the experimental data (a) and the Threshold model (b). * shows significant difference (p < 0.05) between the observed and null data at 50ms using a Permutation test.

## Discussion

To characterize how beta bursts might be involved in action-stopping, we studied three different race model variants; the Interaction model, the Speed-Stop model and the Threshold model. We derived predictions for each model variant on how beta affects behavior in the Stop-signal task. Then we tested these predictions using experimental data and found that the Threshold model best explained the effects of beta bursts in both stopping and going.

While all three models made similar predictions for how the probability of responding is modulated by beta bursts, only the Threshold model could account for the effects seen in Go trials. In Go trials the offset of beta bursts was more likely to occur close to the Go RTs, especially for longer RTs. This similarity between the Threshold model and the experimental data is evidence for a functional role of beta in modulating decision thresholds. Furthermore, the parameters obtained from fitting the Threshold model to participant behavior showed that beta increased Go thresholds more than Stop thresholds, indicating that the effect of beta on stopping can also be indirect by affecting going. Finally, the model correctly predicted the relationship between burst time and action cancellation time, measured as single-trial SSRTs (Hannah *et al.*, 2020), suggesting that for successful stopping the threshold modulation has to occur close and prior to it.

A main finding from our study was that the relationship between the offset of beta bursts and Go RTs showed that beta reflects a short-lived threshold increase, transiently holding the response from execution and thereby helping the Stop-process to win the race. However, as this relation was apparent during Go trials, and not just as a response to the stop cue, it suggests that frontal beta could also be recruited as a proactive mechanism. There is some support for this view from our previous study, in which we found that the beta burst probability increased even in Go trials around the time period when a Stop-signal might have occurred (Jana *et al.*, 2020). Furthermore, for sensorimotor beta there is evidence that it can be proactively recruited (Muralidharan et al., 2019; Soh et al., 2021). Future studies could investigate whether beta increases decision thresholds more generally, or whether this function is adopted by the frontal beta machinery specifically as a preparation to stop if necessary.

Our study also adds more evidence to models involving multiple stages of stopping, such as the pause-then-cancel model (Schmidt and Berke, 2017; Tatz et al., 2021). In the Threshold model beta effectively pauses both decision processes, buying time for the stop process to catch up. This is similar to what we had previously suggested as a potential role for fast stop responses in rat STN neurons (Mallet et al., 2016; Schmidt et al., 2013), which indicates a possible functional connection between sensory responses in basal ganglia neurons and frontal beta. However, in both rodent (Leventhal et al., 2012) and human studies (Jana *et al.*, 2020) there seems to be a longer time gap between the stop cue and the beta burst (120ms in humans) compared to the fast STN responses (~15ms in rats; Schmidt et al., 2013). Therefore, the putative “pause” signal carried by the beta burst here, would instead be in close proximity to the single-trial SSRT measured in the EMG around 160ms. While this could just reflect species differences, it could also be an indication for multiple “pause” systems operating on different, fast and slow, timescales. Importantly, in the Threshold model the beta-mediated pause signal was not triggered by the Stop signal, but instead just occurred randomly. Nevertheless, in the Threshold model we observed that the beta-driven threshold increase occurred close to the SSRT. This was a result of the beta bursts being short-lived, so that the threshold increase only matters if the stop-process is able to catch up with the Go process during the burst. This is exactly the case for beta bursts occurring close to the SSRT because at that time, changes in the threshold matter for the outcome of the race between Go and Stop. While this demonstrates that beta bursts at random time points can lead to temporally specific effects on stopping, it does not preclude that the probability of beta bursts could also be modulated e.g. by sensory events. If our interpretation is correct, a late pause process that occurs close to the SSRT would then overlap in time with any cancelation processes. Therefore, it might be difficult to dissociate them in a standard Stop-signal paradigm.

Even though race models may be considered primarily as phenomenological models addressing behavior, several studies have shown that race models also connect mechanistically to neurophysiology (Hanes and Schall, 1996; Schmidt *et al.*, 2013). In addition, neurophysiological correlates of parameters in rise-to-threshold models have been proposed. For instance, Cavanagh et al. (2011) used drift diffusion models to show that EEG theta oscillations are linked to decision thresholds in a conflict paradigm. Furthermore, the behavioral effects of pharmacological manipulations of striatal dopamine levels could be accounted for by adjusting threshold and accumulation rates (Leventhal et al., 2014). Similarly, in this work we investigated whether some of the functional roles of beta oscillations could be captured in the race model to explain action-stopping. This further supports the wide-applicability of the race model framework to not only account for behavioral data, but to also capture some aspects of the underlying neural processes. However, how beta in the end affects the activity of individual neurons remains a major open question.

Our findings have implications for functions of frontal beta beyond action-stopping. For instance, prefrontal beta has been associated with the executive control of thoughts and memories (Castiglione et al., 2019; Lundqvist et al., 2018; Lundqvist et al., 2016). During thought control, there is increased prefrontal beta in trials, in which participants are successfully preventing the thought from coming to mind (Castiglione *et al.*, 2019). If cognitive inhibition of thought employs mechanisms overlapping with action-stopping, then beta bursts could also reflect a transient threshold increase that helps preventing the thought from reaching consciousness. Furthermore, prefrontal beta has also been shown to play a role in working memory, especially in protecting the current contents of working memory (Lundqvist *et al.*, 2018). A similar mechanism could be at play here where a threshold increase could stop other task-irrelevant stimuli from entering working memory, in line with the classic “status-quo” hypothesis (Engel and Fries, 2010).

In summary, we proposed several models introducing the influence of beta (bursts) into the race framework. We demonstrated that experimental data fitted best to the predictions made by the model that had beta bursts increasing decision thresholds, aiding the stop-process to win the race. Our results provide a clear function role of frontal beta in decision making and the underlying neural mechanisms.

## Acknowledgements

This work was supported by the National Institutes of Health (DA026452), the James S McDonnell Foundation (220020375) and Horizon 2020 Framework Programme (Human Brain Project SGA-3, 945539).

## Supplementary Information

### Beta bursts relationship to going and stopping in Dataset-2

We reproduced our main findings in a second dataset (Dataset-2). First, we validated our beta burst results in this dataset, by looking in a right frontal IC (Fig. S1a) for the percentage of beta bursts (burst %) during successful, failed stops and go trials (Fig. S1b). Our results corroborated with previous studies (Hannah *et al.*, 2020; Jana *et al.*, 2020), where burst % increased during Successful stops in the period of interest (Stop-signal to SSRT) compared to a baseline window prior to stop. Moreover, the burst % in successful stop trials were greater than in go trials. We also replicated our previous results on the relation between beta burst timing and stopping, i.e. the average burst time related to SSRT across participants (Fig. S1c, r = 0.56, p = 0.006, since we did not record EMG in this dataset, we looked at SSRT instead of our EMG metric CancelTime). Next, we looked at the P(Respond) at each burst bin (burst time ± 25ms) and observed the same effect as in Dataset-1. There was a decrease in P(Respond) just before and around SSRT (Fig. S1d, permutation test followed by Bonferroni correction), just as predicted by the threshold model. The joint probability distribution between the Go RTs and burst offset times was also similar to Dataset-1 with more bursts ending closer to the RT, specifically for longer RTs (Fi. S1e). Furthermore, the difference between the Go RTs and burst offsets was skewed towards zero and small positive values for the data compared to a null distribution where bursts were considered to be uniformly distributed for a given RT (Fig. S1f, Kolmogorov-Smirnov test, KSstat = 0.11; p < 0.001)). These results reproduce our main findings from Dataset-1 and support that beta bursts modulate decision thresholds.

**Fig. S1.**
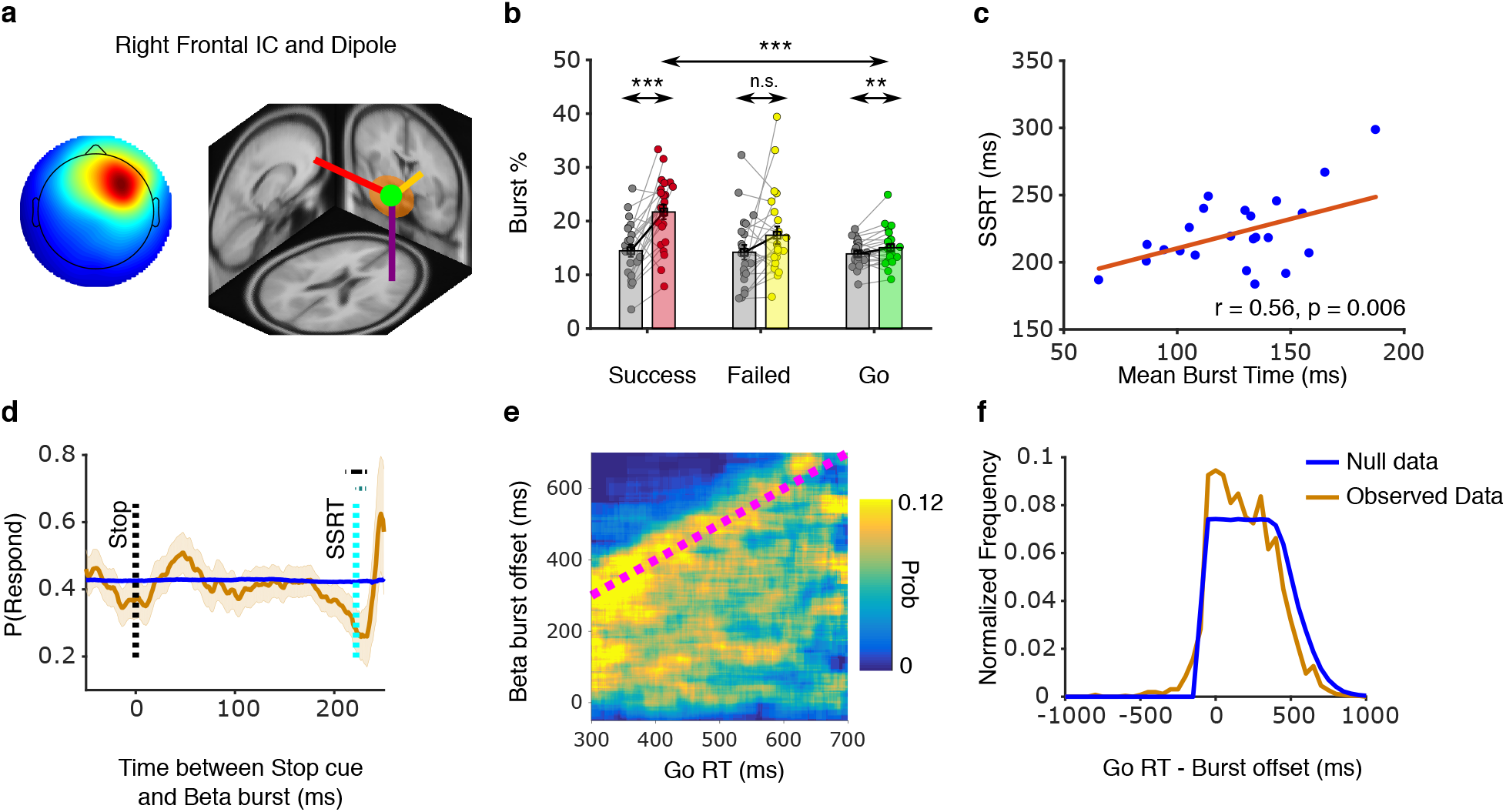
Beta bursts and behavior for Dataset-2. a) Average right frontal independent component and average dipole location (green dot) of the right frontal ICs, show a right lateralized frontal topography for the selected ICs. b) Burst % in successful stop (red bar), failed stop (yellow bar) and go trials (green bar) in the period between Stop-signal and SSRT compared to a baseline prior to stop (corresponding grey bar). The burst % increases in successful stop trials compared to its baseline and also this increase is significantly higher compared to that of go trials. c) Linear relationship between burst time and SSRT across participants. d) Probability of response as a function of burst-time bins in experimental data for Dataset-2. The probability estimate is for each burst-bin which represents bursts in the time window burst time±25ms. The solid black line and pale blue line above denote the uncorrected and Bonferroni corrected p values respectively, showing time periods of significant decreases in probability of response w.r.t to the null distribution (solid blue line, obtained by shuffling the labels of success and failures) using a permutation test. e) The joint probability distribution between the burst offset times and Go RT bins. The burst offsets are defined in relation to the Go cue. Like in panel (d) the probability estimate at each pixel/bin is obtained from reaction times and burst offset times in a time window ±25ms around the particular RT and burst offset. The diagonal dotted magenta line represents the line where RT is equal to burst offset. f) The normalized distribution of the difference between Go RTs and burst offset time, compared to a null-distribution (solid blue line).

### Varying beta burst duration in the Threshold model

For Dataset-1, we fitted the parameters for β_dur_ = 100ms and simulated the Threshold model using different durations of a burst (β_dur_ = 125ms and 150ms). The pattern of modulation of the probability of response was similar across the different burst durations, which decreased close to SSRT, although the decrease was a bit pronounced for longer β_dur_ (Fig. S2a). The relationship between the simulated Go RTs and burst offset times were also similar across all the three burst durations, with an increase in the probability of bursts ending close to the RTs (Fig. S2b).

**Fig. S2.**
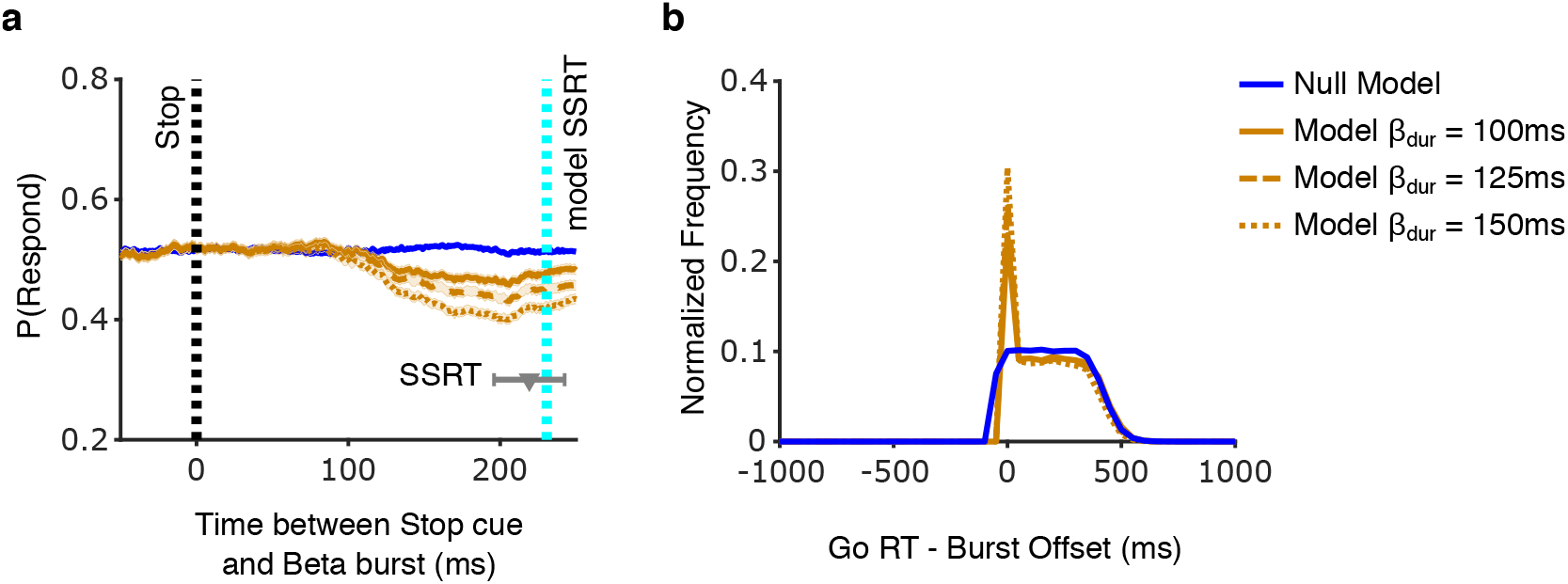
Varying burst durations in the Threshold model. a) Probability of response across burst bins for different burst durations (100ms, 125ms and 150ms). b) Normalized distribution of the difference between simulated Go RTs and burst offset times for different burst durations.

### Threshold Model parameters for different number of trials with bursts

The model parameters of the beta bursts (Fig. 2g-i) were obtained from simulations in which beta bursts occurred in 20% of the trials, similar to the experimental data. To see whether the percentage of trials with beta bursts affects the model fit and parameters, we also performed a model fit in which beta bursts occurred on every trial (Fig. S3). The model reaction time distributions and inhibition functions were similar irrespective of the percentage of trials with beta, and both provided a good fit with the experimental behavioral data (Fig. S3a-c). Furthermore, the model with beta bursts in every trial exhibited the same pattern of the beta burst parameters (Fig. S3d-f) as the 20% model (Fig. 2g-i).

**Fig. S3.**
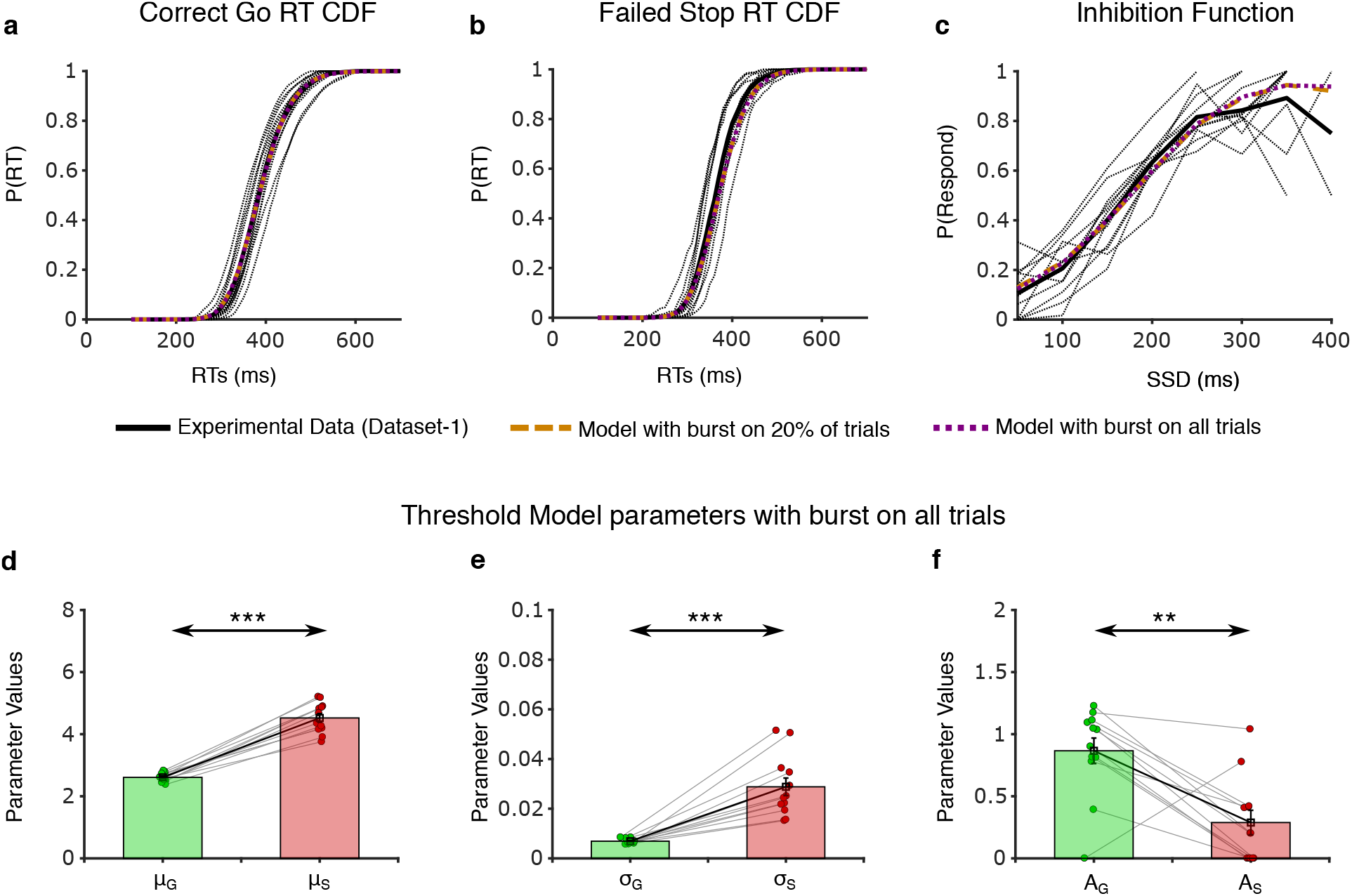
Threshold Model Parameters with different bursts percentage. a-c) Comparison of reaction time distributions and inhibition functions between experimental and model data. The solid black line shows the average of the experimental data and the dotted black lines show individual participants from Dataset-1. Shown are Correct Go RT CDFs (a), Failed Stop RT CDFs (b), and Inhibition function (c) for both model scenarios, i.e. 20% of trials having a burst (dashed brown line) and all trials having a burst (dotted purple line). d-f) Fitted parameters for the model with all trials having a burst, i.e. mean slope μ (d), slope standard deviation σ (e), and threshold modulation A (f) for the go and stop process.

### Threshold model: Relationship between Go RTs and burst offsets with noise

The effect of threshold modulation during the Go trials in the experimental data was observed as a diagonal pattern in the beta burst offset and Go RT probability distribution (Fig. 2b). However, the pattern of modulation seen in the threshold model was much sharper than in the experimental data (Fig. 2e, f). We suspected that noise in estimation of the burst offset could contribute to the wider diagonal pattern seen in the experimental data. To test this idea, we added jittered the burst offset times in the model by adding a normally distributed random number to the burst offset time in each trial. We examined two scenarios, with the standard deviation of the noise being either 25ms or 50ms. We then recomputed the joint probability distribution with these two different noise factors and observed that indeed the sharp diagonal stripe became broader upon adding noise (Fig. S4b, c, e, and f), increasing the similarity between the model and the empirical data. This supports our idea that the differences between the model and the empirical data with respect to the width of the pattern in the joint probability distribution can indeed be explained by the fact that the offset of a beta oscillation can only be estimated in the empirical data.

**Fig. S4.**
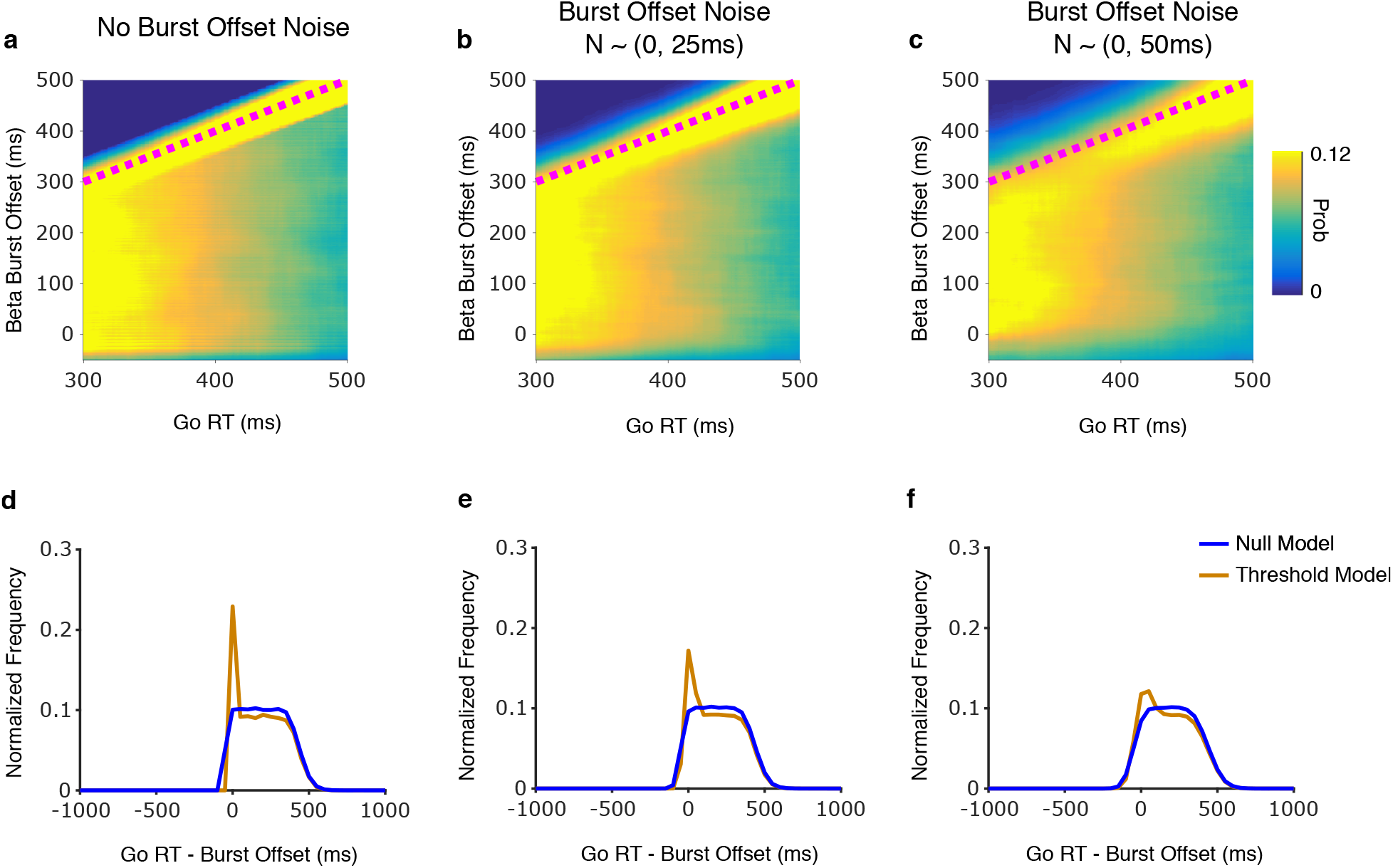
The relation between **Go RTs and Burst Offsets in the threshold model with noise**. a-c) The joint probability distribution between the burst offset times and Go RTs for different noise ranges added to the burst offset times; no noise (a), uniformly random noise from −50 to 50ms (b) and −100 to 100ms (c). d-f) The normalized distribution of the difference between Go RTs and burst offset time, compared to a null-distribution (solid blue line) for different noise factors; no noise (d), uniformly random noise from −50 to 50ms (e) and −100 to 100ms (f). The diagonal modulation becomes broader with increasing noise range.

